# The Network Basis for the Structural Thermostability and the Functional Thermoactivity of Aldolase B

**DOI:** 10.1101/2022.10.20.513014

**Authors:** Guangyu Wang

**Affiliations:** Department of Physiology and Membrane Biology, University of California School of Medicine, Davis, CA, USA; Department of Drug Research and Development, Institute of Biophysical Medico-chemistry, Reno, NV, USA

**Keywords:** grid thermodynamics, enzyme, non-covalent interaction, systemic thermal instability, temperature sensitivity, thermostability, thermoactivity, threshold

## Abstract

Thermostability is important for thermoactivity of proteins including enzymes. However, it is still challenging to pinpoint the specific structural factors for different temperature thresholds to initiate their specific structural and functional perturbations. Here, graph theory was used to investigate how the temperature-dependent noncovalent interactions as identified in the structures of aldolase B and its prevalent A149P mutant could form a systematic fluidic grid-like mesh network with topological grids to regulate the structural thermostability and the functional thermoactivity. The results showed that the biggest grid may determine the temperature thresholds for the changes in their secondary and tertiary structures and specific catalytic activities. Further, a highly conserved thermostable grid may serve as an anchor to secure the flexible active site to achieve the specific thermoactivity. Finally, higher grid-based systematic thermal instability may disfavor the thermoactivity. Thus, this computational study may provide critical clues for the structural thermostability and the functional thermoactivity of proteins including enzymes.

## 1. INTRODUCTION

Thermostability of proteins including enzymes is a keystone for their thermoactivity. Usually, thermostability is determined by a melting temperature (T_m_) at which the unfolding of the secondary structure causes protein denaturation. However, when a missense mutation renders protein a temperature threshold (Tth) for inactivation, a change in the tertiary or quaternary structure may be involved even if the overall secondary structure is intact [1–2]. A good example is aldolase B, which is critical for gluconeogenesis and fructose metabolism. The most common mutant, A149P, is responsible for about 55% of hereditary fructose intolerance (HFI) alleles worldwide [3]. HFI causes excessive accumulation of Fru 1-P in tissues expressing aldolase B. Several hazardous metabolic effects induce liver and kidney injury if untreated [4–5]. Therefore, early intervention or the dietary exclusion of noxious foods containing fructose, sucrose, and sorbitol is necessary for the HFI management.

Human aldolase B with or without the A149P substitution have been studied structurally and functionally [1–2]. Regarding the specific activity toward Fru-1,6-P_2_ and Fru-1-P, the wild-type construct exhibits the low catalytic activity above 40 °C due to a loss in the tertiary structure. It has no activity at 50 °C because the secondary structure has been affected. In contrast, its A149P mutant decreases the specific activity for Fru-1-P cleavage to 16% of the wild-type level at 10 °C. Its activity further declines above 15 °C. When the tertiary structure is lost at 30 °C, its specific activity is as low as 0.5% of the wild-type level. It has no activity at 40 °C as a result of perturbations in the secondary and tertiary structures [1]. In addition, although wild-type aldolase B is tetrameric, the A149P mutant has dimeric species in solution from 4 °C to 30 °C and monomeric species above 30 °C [1]. Further investigations indicated that some structural rearrangements result in the low activity. They include disordered loop regions from residues 148-159 to residues 110-129, 189-199, and 235-242. Notably, the critical E189-R148 salt bridge at the active site is broken at 18 °C [2]. Thus, these mutation-induced structural perturbations account for the mutation-evoked HFI very well. However, until now the specific structural factors or motifs for different T_th_ parameters to affect the structural thermostability and the functional thermo-activity are still missing.

A key clue comes from a DNA hairpin thermal biosensor. When the loop size is not beyond 30 bases, its T_m_ is primarily controlled by the loop size and H-bonds in the stem and thus can be employed to monitor a change in the environmental temperature with a high sensitivity. The bigger hairpin size and the less H-bonds in the stem usually have a lower T_th_ [6]. Its melting curve is also similar to the temperature dependent activity of aldolase B and its A149P mutant because both have a characteristic sigmoid slope [1, 6]. As both a grid in a grid-like non-covalent interaction mesh network and a single DNA hairpin have a common topological circle in general term and the dissociation of this network grid by heat is analogous to DNA hairpin melting, it is exciting to hypothesis that aldolase B and its A149P mutant use a fluidic system of temperature-dependent grid-like non-covalently interacting mesh networks with topological grids to keep the 3D structure of aldolase B for the different structural thermostability and functional thermoactivity.

In this computational study, graph theory was exploited as an *ab initio* method to examine the hypothesis by carefully deciphering each grid in the individual grid-like non-covalently interacting mesh networks as identified in the crystal structures of wild-type aldolase B at room temperature (293 K) and its mutant A149P at 277 K and 291 K [2, 7–8]. Melting temperature (T_m_) of the biggest grid and grid-based systematic thermal instability (Ti) were then similarly derived on the basis of the structural and functional data of DNA hairpins to pinpoint the structural factor or motif as the ground for the structural thermostability and the functional thermoactivity.

It has been reported that the temperature threshold above 34 °C can be increased more than 20 degrees when the H-bonded G-C base-pairs double from 2 to 4 or the loop length in a DNA hairpin declines from 20 to 10 A’s [6]. Accordingly, a grid size (S) and grid size-controlled equivalent H-bonds in the network grid may play a critical role in the melting temperaure. In this computational study, once the grid size was defined by graph theory as the minimal number of the total side chains of residues in protein that did not involve any non-covalent interctions in a grid, the biggest grid along the single polypeptide chain could then be identified. If the biggest grid undergoes the single-rate-limiting-step melting reaction like DNA hairpins for a change in the tertiary or secondary structure or the specific catalytic activity in aldolase B, the calculated melting temperature (T_m_) of the biggest grid should be comparable to the experimental T_m_ or threshold (Tth).

On the other hand, smaller loops or more H-bonds in the stem have been reported to enhance the thermostability of a DNA hairpin [6]. Hence, if the whole grid-like non-covalent interaction mesh network along the single polypeptide chain rearranges along with the melting of the biggest grid, systemic thermal instability (T_i_), once defined as a ratio of the total grid sizes to the total non-covalent interactions along the same single polypeptide chain, could be an important energetic reference for the structural thermostability and the functional thermoactivity.

Based on two above lines of comparative parameters, it was found that the grid-based systematic thermal instability (T_i_) was significantly increased by the A149P mutation but different between paired monomers. Since several calculated T_m_ values of the biggest grids were found consistent with the measured T_th_ values for the specific structural and functional changes in either aldolase B or its A149P mutant, such a grid thermodynamic model may be a powerful tool to predict the temperature thresholds for the changes in the structural thermostability and the functional thermoactivity of proteins including enzymes.

## 2. MATERIALS AND METHODS

### 2.1. Data Mining Resources

In this computational study, three X-ray crystallographic structures of the isoenzyme aldolase B in the presence of (NH_4_)_2_SO_4_ were analyzed with graph theory to reveal the roles of the biggest grids in regulating the T_th_ parameters for changes in their tertiary or secondary structures and thermal activity. One was the human liver isozyme of fructose-1,6-bisphosphate aldolase B at room temperature (293 K) (PDB ID, 1QO5, model resolution= 2.5 Å) [9]; two others were its A149P mutant at 277 K (PDB ID, 1XDL, model resolution= 3.0 Å) and 291 K (PDB ID, 1XDM, model resolution= 3.0 Å) [2]. Because the disordered loop region in monomer C or W of the A149P mutant at 277 K or 291 K has affected a critical E189-R148 salt bridge in the active-site, only paired monomers A and B in the active aldolase-A149P tetramer and paired monomers A and D in the wild-type active aldolase tetramer were employed for the grid thermodynamic modeling. On the other hand, paired monomers Z and Y, which are similar to monomer A and B, respectively, were not further analyzed [2].

### 2.2. Standards for Non-covalent Interactions

Structure visualization software, UCSF Chimera, was used to identify stereo- or regio-selective inter-domain diagonal and intra-domain lateral non-covalent interactions in aldolase B and its mutant A149P, and to test their potential roles in forming topological grids with minimal sizes to control the T_th_ for the changes in the tertiary or secondary structures and specific activity of wild-type aldolase B and its A149P mutant. These non-covalent interactions included salt-bridges, CH/π-π interactions and H-bonds along the single polypeptide chain from T8 to S352.

The standard definition of noncovalent interactions was employed as a filter to secure that results could be reproduced with a high sensitivity. An H-bond was considered effective when the angle donor-hydrogen-acceptor was within the cut-off of 60°, and when the hydrogen-acceptor distance was within 2.5 Å and the maximal distance was 3.9 Å between a donor and an acceptor. A salt bridge was considered valid if the distance between any of the oxygen atoms of acidic residues and the nitrogen atoms of basic residues were within the cut-off distance of 3.2-4 Å in at least *one pair. When* the geometry was acceptable, a salt bridge was also counted as a hydrogen bonding pair. The face-to-face π-π stacking of two aromatic rings was considered attractive once the separation between the π-π planes was ~3.35-4.4 Å, which is close to at least twice the estimated van der Waals radius of carbon (1.7 Å). The edge-to-face π-π interaction of two aromatic rings was also considered attractive when the cut-off distance between two aromatic centers was within 4-6.5 Å. The short effective CH/π distances was 2.65-3.01 Å between aromatic groups, and 2.75-2.89 Å between CH3 and an aromatic ring.

### 2.3 Preparation of Topological Grid Maps by Using Graph Theory

After non-covalent interactions were scanned along the single polypeptide chain from T8 to S352, graph theory was used to define a grid as the smallest topological circle that had the minimal size to control a non-covalent interaction in the grid. All the grids with minimal sizes were geometrically mapped for aldolase B at room temperature (293 K) and its A149P mutant at 277 K and 291 K. The primary amino acid sequence line from T8 to S352 was marked in black. An amino acid side chain involving a non-covalent interaction along the single polypeptide chain was represented as a vertex (*v*) or a node and marked with an arrow in different colors. The same kind of non-covalent interactions was marked in the same color. Any link between two vertices was represented an edge in a biochemical network. A topological grid was formed between two vertices *i* and *j* (v_*i*_ and v_*j*_), if and only if there was a path from v_*i*_ to v_*j*_ and a return path from v_*j*_ to v_*i*_. The grid size (S) was definded as the minimal number of the total side chains of residues that did not participate in any non-covalent interctions in a grid. For a given grid-like biochemical reaction mesh network, the grid size between v_*i*_ and v_*j*_ was the sum of the shortest return path distance from node *j* to *i* because the direct shortest path distance from node *i* to *j* was zero when a non-covalent interaction existed between them. The simplest way to find the grid size was to identify the shortest return path between two vertices *i* and *j* (v_*i*_ and v_*j*_) by using the Floyd-Warshall algorithm [10]. For example, in the biochemical reaction network of Fig.1A, a direct path length from K12 and Y222 was zero because there was a CH-π interaction between them. However, there was another shortest return path from Y222 to T226, Q178, Y173, W147, D143, K138, E14 and back to K12 via two H-bonds and a salt bridge and a π–π interaction in this grid. Because the total number of residues that did not engage in non-covalent interactions in the grid was 11, the grid size was 11. After each non-covalent interaction was tracked by a grid size and the uncommon sizes were marked in black, a grid with an x-residue or atom size was denoted as Gridx, and the total non-covalent interactions and grid sizes along the single polypeptide chain were shown in black and blue circles beside the network map, respectively.

**Figure 1.**
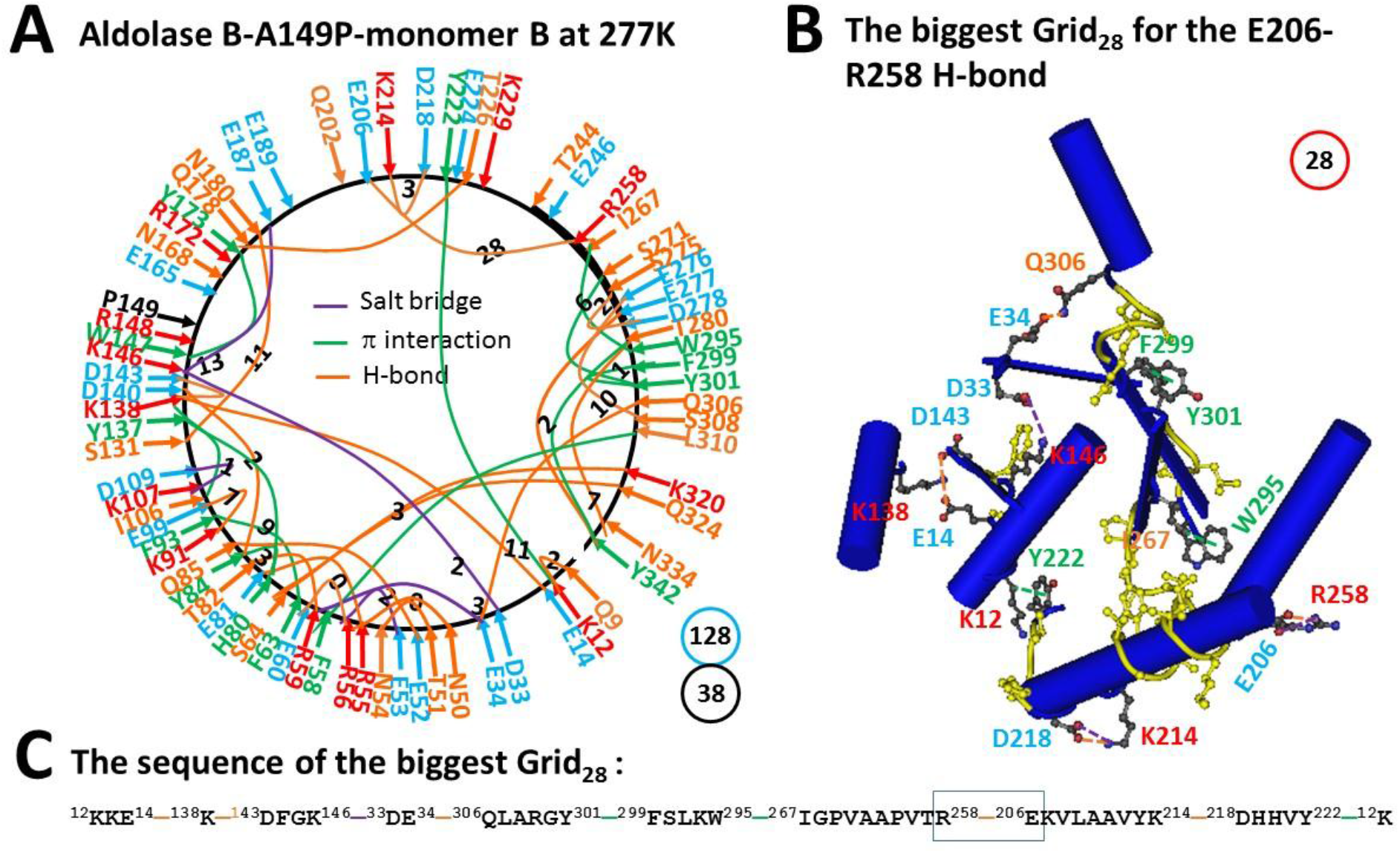
The grid-like non-covalently interacting mesh network along the single polypeptide chain of ligand-free monomer B in the aldolase B-A149P tetramer at 277 K. **A.** The topological grids in the systemic fluidic grid-like mesh network. The X-ray crystallographic structure of monomer A in aldolase B-A149P at 277 K (PDB ID, 1XDL) was used for the model [2]. Salt bridges, π interactions, and H-bonds between pairing amino acid side chains along the single polypeptide chain from T8 to Y342 are marked in purple, green, and orange, respectively. The grid sizes required to control the relevant non-covalent interactions were calculated with graph theory and labeled in black. The total grid sizes and grid size-controlled non-covalent interactions along the single polypeptide chain are shown in the blue and black circles, respectively. **B,** The structure of the biggest Grid_28_. The grid size is shown in a red circle. **C,** The sequence of the biggest Grid_28_ to control the E206-R258 H-bond in a blue box.

### 2.4 Equation

When a non-covalent interaction in a grid is broken at a temperature, that temperature was defined as a melting one. If both a DNA hairpin and the biggest grid melts in a similarly manner and the melting is rate-limiting for the primary structural and functional changes, the T_th_ for the changes in the tertiary or secondary structures or the thermal activity of proteins was calculated from the T_m_ of the biggest grid along the single polypeptide chain using the following equation [6]:

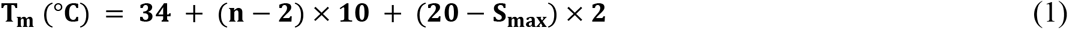

where, n is the total number of the grid size-controlled non-covalent interactions equivalent to H-bonds in the biggest grid, and S_max_ is the size of the biggest grid.

In either state, grid-based systemic thermal instability (T_i_) along the single polypeptide chain was defined using the following equation:

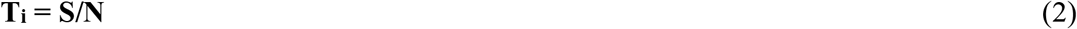

where, S is the total grid sizes along the single polypeptide chain of one subunit in a functional state; N is the total non-covalent interactions along the same single polypeptide chain of one subunit in the same functional state.

### 2.5 Ethics Statement

This computational study did not involve any human participants or human subjects research or retrospective study of medical records or archived samples.

## 3 RESULTS

### 3.1. The Biggest Grid_28_ in Monomer B of Aldolase B-A149P at 277 K Had a Predicted T_m_ 18 °C

The ligand-free aldolase B-A149P tetramer has paired monomers A and B which are linked by two swapping salt bridges between E224 from one monomer and R258 from the other at their interface [2]. Although both monomers had the same primary sequence, they had some different non-covalent interactions between amino acid side chains in each subunit to produce distinct grids [2].

First, eighteen H-bonds were present between various hydrophilic residues along the single polypeptide chain from T8 to Y432. The H-bonding pairs included Q9-K12, E14-K138, E34-Q306, N50/E81-R55, T51-N54, E52-Q85, R56-E60, R59-T82, S64-K320/Q324, Y84-D140, K91-E99, S131-N180, K138-D143, Q178-T226, E206-R258, K214-D218, S275-D278, E276-S308, E277-Y342 and T280-N334 (Fig. 1A).

Second, eleven π interactions were formed between aromatic residues and nearby residues. For example, CH-π interacting pairs had K12-Y222, F58-L310, I106-Y137, I267-W295, S271-Y301 and T280-Y432 while π–π interacting pairs covered F63-F93, H80-Y84/Y137, W147-Y173 and F299-Y301 (Fig. 1A).

Third, nine salt bridges were also observed between several charged pairs. They included D33-K146, E34-R59, E53-R56, K107-D109, K138-D143, and K146-E187 (Fig. 1A).

Taken together, thirty-eight non-covalent interactions generated multiple grids. Their sizes were in a range from 0 to 28 and had a total number 128 (Table 1). Thus, the systematic thermal instability was 3.37 (Table 1). Of special note, the biggest Grid_28_ had a 28-residue size via the shortest path from K12 to E14, K138, D143, K146, D33, E34, Q306, Y301, F299, W295, I267, R258, E206, K214, D218, Y222 and back to K12. Because this biggest Grid_28_ had 2.0 equivalent H-bonds to seal the grid for the E206-R258 salt bridge, its T_m_ was calculated as about 18 °C (Fig. 1B-C, Table 1).

**Table 1.** New parameters of the ligand-free aldolase B based on the grid thermodynamic model.

| Construct | Wild-type |  | A149P |  |  |  |
| --- | --- | --- | --- | --- | --- | --- |
| PDB ID | 1QO5 |  | 1XDL |  | 1XDM |  |
| Sampling temperature | Room temperature (293K) |  | 277 K |  | 291 K |  |
| Monomer in a tetramer | A | D | A | B | A | B |
| Name of the biggest Grid | Grid<sub>11</sub> | Grid<sub>14</sub> | Grid<sub>27</sub> | Grid<sub>28</sub> | Grid<sub>14</sub> | Grid<sub>25</sub> |
| Biggest grid size (S<sub>max</sub>), a.a | 11 | 14 | 27 | 28 | 14 | 25 |
| Equivalent H-bonds in S<sub>max</sub> | 2 | 1.5 | 1.5 | 2.0 | 1.5 | 2.5 |
| Total non-covalent interactions (N) | 49 | 47 | 37 | 38 | 37 | 39 |
| Total grid sizes (S), a.a. | 140 | 139 | 135 | 128 | 126 | 147 |
| Systematic thermal instability | 2.86 | 2.96 | 3.65 | 3.37 | 3.41 | 3.77 |
| Calculated T<sub>m</sub> | 52 °C | 41 °C | 15 °C | 18 °C | 41 °C | 29 °C |
| Measured T<sub>th</sub> for tertiary structure | 50 °C | 40 °C |  |  | 40 °C | 30 °C |
| Measured T<sub>th</sub> for the activity | 50 °C | 40 °C | 15 °C | 15 °C | 40 °C |  |
| Ref. for T<sub>th</sub> | [1-2] | [1-2] | [2] | [2] | [1-2] | [1] |

### 3.2. The Melting of the Biggest Grid_28_ in Monomer B of Aldolase B-A149P at 291 K produced the Biggest Grid_25_ with a Calculated T_m_ 29 °C

As predicted, the biggest Grid_28_ in monomer B of aldolase B-A149P did melt at 18 °C when the E205-R258 H-bond was disrupted at 291 K (Table 1). Meanwhile, several changes in non-covalent interactions of this monomer were followed.

First, when H-bonding pairs S131-N180 and Q178-T226 were replaced with those K133-Y137, N166-E189, Q202-E206 and T244-E246 pairs, T280 H-bonded with R330. After the E52-Q85 H-bond was broken, S35 H-bonded with T38, and the Q9-K12 H-bond moved to the T8-Q11 one (Fig.2A).

**Figure 2.**
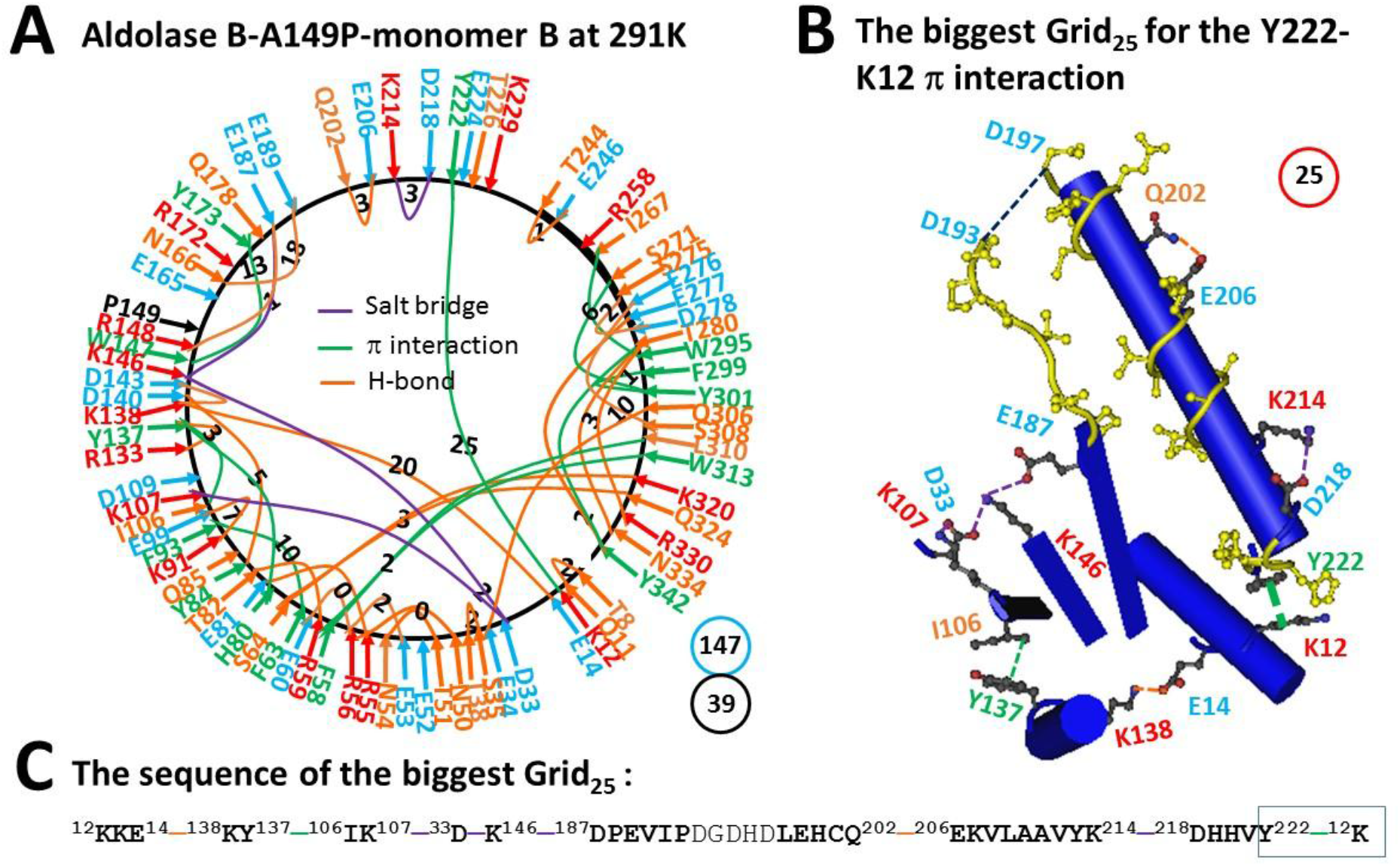
The grid-like non-covalently interacting mesh network along the single polypeptide chain of ligand-free monomer B in the aldolase B-A149P tetramer 291 K. **A.** The topological grids in the systemic fluidic grid-like mesh network. The X-ray crystallographic structure of monomer A in aldolase B-A149P at 291 K (PDB ID, 1XDM) was used for the model [2]. Salt bridges, π interactions, and H-bonds between pairing amino acid side chains along the single polypeptide chain from T8 to Y342 are marked in purple, green, and orange, respectively. The grid sizes required to control the relevant non-covalent interactions were calculated with graph theory and labeled in black. The total grid sizes and grid size-controlled non-covalent interactions along the single polypeptide chain are shown in the blue and black circles, respectively. **B,** The structure of the biggest Grid_25_. The grid size is shown in a red circle. **C,** The sequence of the biggest Grid_25_ to control the Y222-K12 CH-π interaction in a blue box.

Second, when E187 also H-bonded with R148, the R56-E53 and K138-D143 salt bridges became H-bonds but the K214-D218 H-bond changed to a salt bridge. Further, when the E34-R56 and K107-D109 salt-bridges were disconnected, a new D33-K107 salt bridge emerged (Fig. 2A).

Third, with the H80-Y84 π–π interaction being disrupted, a new F58-W313 one was added (Fig. 2A). As a result, although the total non-covalent interactions along the single polypeptide chain was slightly increased from 38 to 39, the total grid sizes significantly increased from 128 to 147 (Figs 1A, 2A). Thus, the systematic thermal instability was increased from 3.37 to 3.77 (Table 1). After the biggest Grid_28_ melted at 291 K, the biggest Grid_25_ was present with a 25-residue size to control the strong K12-Y222 CH-π interaction. It started from K12, covered E14, Y137, I106, K107, D33, K146, D187, Q202, E206, K214 and D218, and ended with Y222 (Fig. 2B-C). When 2.5 equivalent H-bonds were taken into account, a T_m_ 29 °C was generated (Fig. 2B, Table 1).

### 3.3. The Biggest Grid_27_ in Monomer A of Aldolase B-A149P at 277 K Had a Predicted T_m_ 15 °C

In contrast to monomer B, monomers A of aldolase B-A149P at 277K exhibited different non-covalent interactions and related grids. When the D33-K146 salt bridge shifted to the D33-K107 one near the mutation site, the Q9-K12 H-bond also moved to the T8-Q11 one, the E53-R56 salt bridge changed to an H-bond, and the T51-N54 and E52-Q85 and K91-E99 H-bonds were disconnected. On the other hand, when the new K148-E189 salt bridge and the new E165-R172 H-bond appeared, the Q178-T226 H-bond changed to an E187-K229 salt bridge, the K214-D218 H-bond became a salt bridge, and the Q202-E206 and T244-E246 H-bonds were added but the I267-W295 CH-π interaction was broken. In this case, the total grid sizes raised up to 135 but the total non-covalent interactions slightly decreased to 37 (Fig. 3A). Hence, the systematic thermal instability was 3.65 (Table 1). Of special note, the biggest Grid_27_ had a 27-residue size via the shortest path from K12 to E14, K138, Y137, I106, K107, D33, E34, Q306, Y301, S271, R258, E206, K214, D218, Y222 and back to K12 (Fig. 3B-C). When 1.5 equivalent H-bond sealed the biggest Grid_27_ to control the E206-R258 salt bridge, the calculated T_m_ was 15 °C (Fig. 3B-C, Table 1).

**Figure 3.**
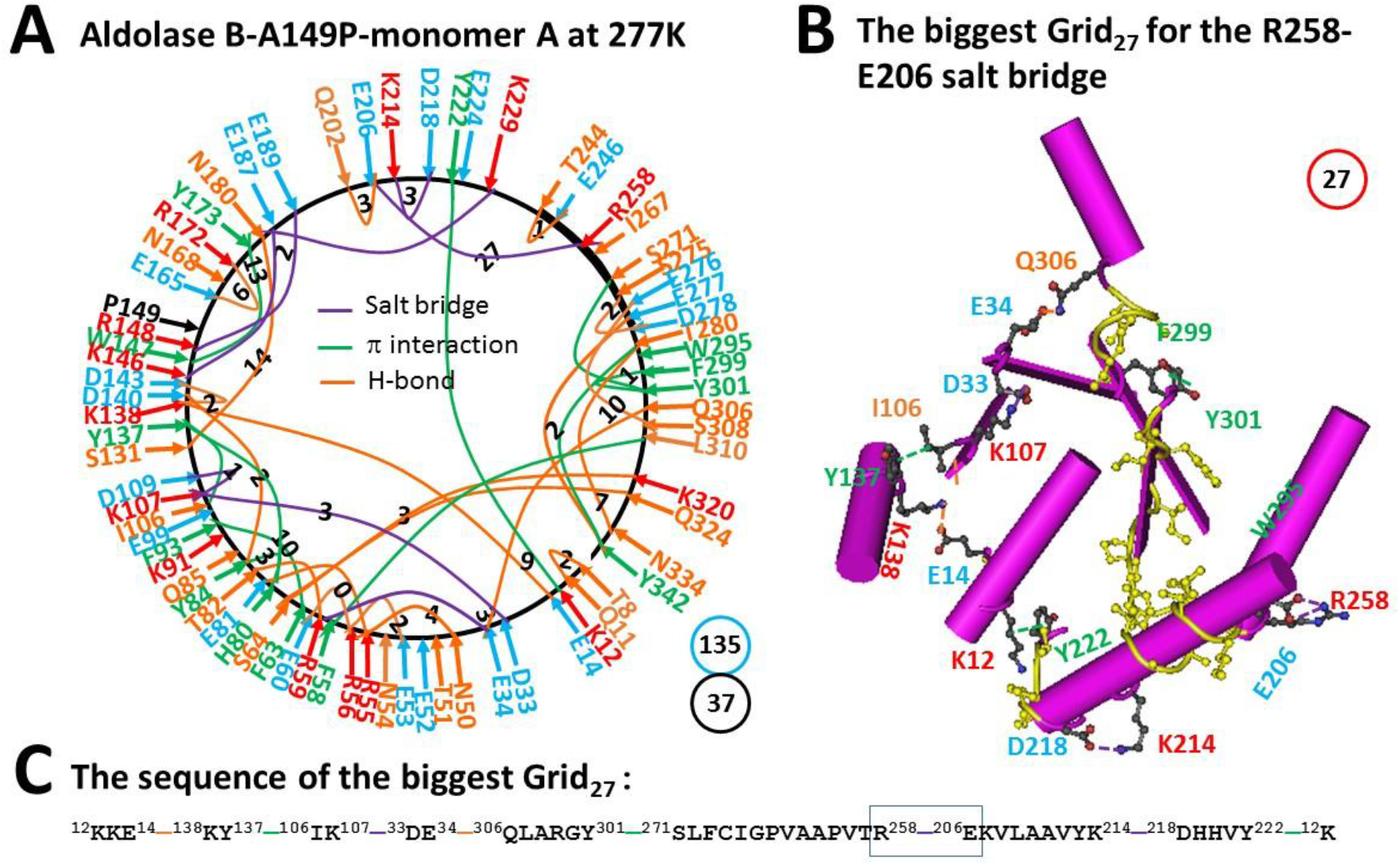
The grid-like non-covalently interacting mesh network along the single polypeptide chain of ligand-free monomer A in the aldolase B-A149P tetramer at 277 K. **A.** The topological grids in the systemic fluidic grid-like mesh network. The X-ray crystallographic structure of monomer B in aldolase B-A149P at 277 K (PDB ID, 1XDL) was used for the model [2]. Salt bridges, π interactions, and H-bonds between pairing amino acid side chains along the single polypeptide chain from T8 to Y342 are marked in purple, green, and orange, respectively. The grid sizes required to control the relevant non-covalent interactions were calculated with graph theory and labeled in black. The total grid sizes and grid size-controlled non-covalent interactions along the single polypeptide chain are shown in the blue and black circles, respectively. **B,** The structure of the biggest Grid_27_. The grid size is shown in a red circle. **C,** The sequence of the biggest Grid_27_ to control the E206-R258 salt bridge in a blue box.

### 3.4. The Melting of the Biggest Grid_27_ in Monomer A of Aldolase B-A149P at 291 K produced the Biggest Grid_14_ with a Calculated T_m_ 41 °C

Unlike monomer B, the E206-R258 salt bridge was not disrupted when temperature increased from 4 °C to 18 °C. Instead, a nearby Y213-P261 CH-π interaction was added so that a grid size declined from 27 to 8 residues to keep the E206-R258 salt bridge. As a result, when the E187-K229 salt bridge was substituted by the Q178-T226 H-bond, a H196-H200 π–π interaction emerged but the S131-N180 H-bond and the E189-K148 and E165-R172 salt bridges were disconnected. When the D109-K107-D33 salt bridges were broken, the K91-E99 and Q85-E52 H-bonds were reinstated together with a new R41-N42 H-bond. On the other hand, when the new K229-S300 and S271-R303 H-bonds were added, the S64-K320 H-bond was disrupted so that the biggest Grid_14_ was born. It had a 14-residue size to control the S64-Q423 H-bond. This grid had the shortest path from S64 to Q324, W313, F58, R56, E60 and back to S64. As 1.5 equivalent H-bonds closed the grid, T_m_ was calculated as about 41 °C. In the meanwhile, the total grid sizes lowered down from 146 to 126 (Fig.4). Thus, along with the increase in the systematic thermal instability of monomer B, monomer A exhibited a compensatory decrease in this parameter from 3.65 to 3.41 at elevated temperature (Table 1).

**Figure 4.**
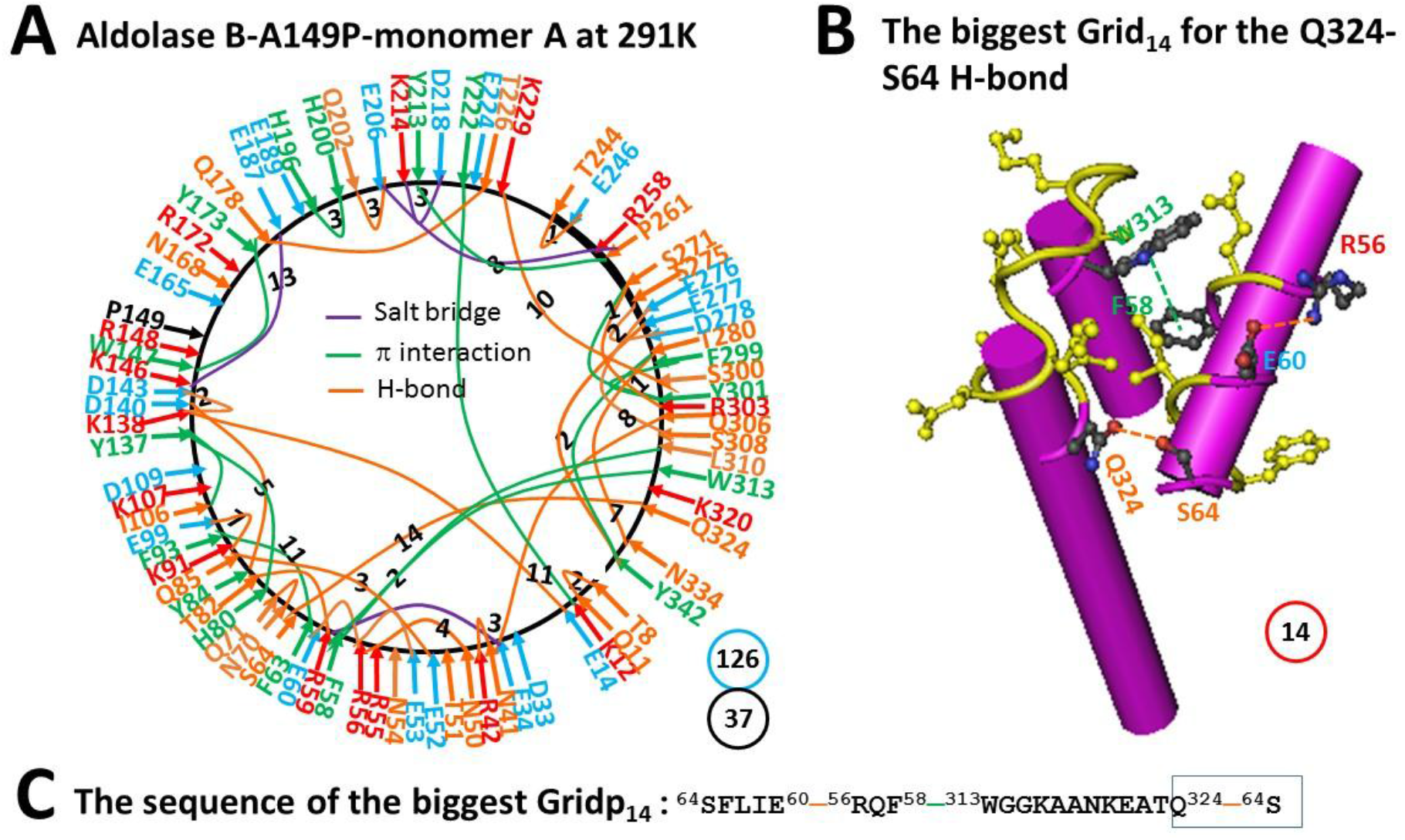
The grid-like non-covalently interacting mesh network along the single polypeptide chain of ligand-free monomer A in the aldolase B-A149P tetramer at 291 K. **A.** The topological grids in the systemic fluidic grid-like mesh network. The X-ray crystallographic structure of monomer A in aldolase B-A149P at 291 K (PDB ID, 1XDM) was used for the model [2]. Salt bridges, π interactions, and H-bonds between pairing amino acid side chains along the single polypeptide chain from T8 to Y342 are marked in purple, green, and orange, respectively. The grid sizes required to control the relevant non-covalent interactions were calculated with graph theory and labeled in black. The total grid sizes and grid size-controlled non-covalent interactions along the single polypeptide chain are shown in the blue and black circles, respectively. **B,** The structure of the biggest Grid_14_. The grid size is shown in a red circle. **C,** The sequence of the biggest Grid_14_ to control the S64-Q324 H-bond in a blue box.

### 3.5. The Same Biggest Grid_14_ with the Same T_m_ 41 °C Appeared in Monomer D of Wild-type Aldolase B at room temperature (293 K)

Since the biggest grids in monomers A and B of aldolase B-A149P did melt at or above predicted T_m_ values (Figs 1-4, Table 1), it is reasonable to predict T_m_ values of wild-type aldolase B by using the biggest grids. Given that monomers D and A in wild type aldolase B had the same dimeric interface as monomers B and A in the A149P mutant [2, 9], the calculations were focused on monomers D and A.

The network grid map of monomer D in wild type aldolase B at room temperature (293 K) was much more like that of monomer A in the A149P mutant at 277 K. However, without the mutation, the E165-R172 salt bridge and the S131-N180 H-bond were absent. As a result, when the E187-K229 salt bridge was broken, the Q178-T226 and D197-K241 H-bonds and the I153-H196-H200 π interactions appeared. When K229 H-bonded with S300, the I267-W295 CH-π interaction was present again with the S275-D278 H-bond moving to the E277-H344 one. On the other hand, when the K146-D33 and E121-T118 and S88-E60 and Q85-E52 H-bonds were introduced, the D109-K107 salt bridge shifted to the E99-K91 one, and the R59-E34 and R56-R53 salt bridges became H-bonds. Taken together, the S64-K320 H-bond was disrupted but the F58-W313 π–π interaction appeared again. In this case, the total grid sizes increased from 135 to 139 along with an increase in the total non-covalent interactions from 37 to 47 (Fig.5A). Thus, the systematic thermal instability was 2.96 for the wild type isoenzyme (Table 1). When 1.5 equivalent H-bonds sealed the same biggest Grid_14_ as found in monomer A of the A149P mutant at 291 K, the calculated T_m_ was also about 41 °C (Fig. 5B-C, Table 1).

**Figure 5.**
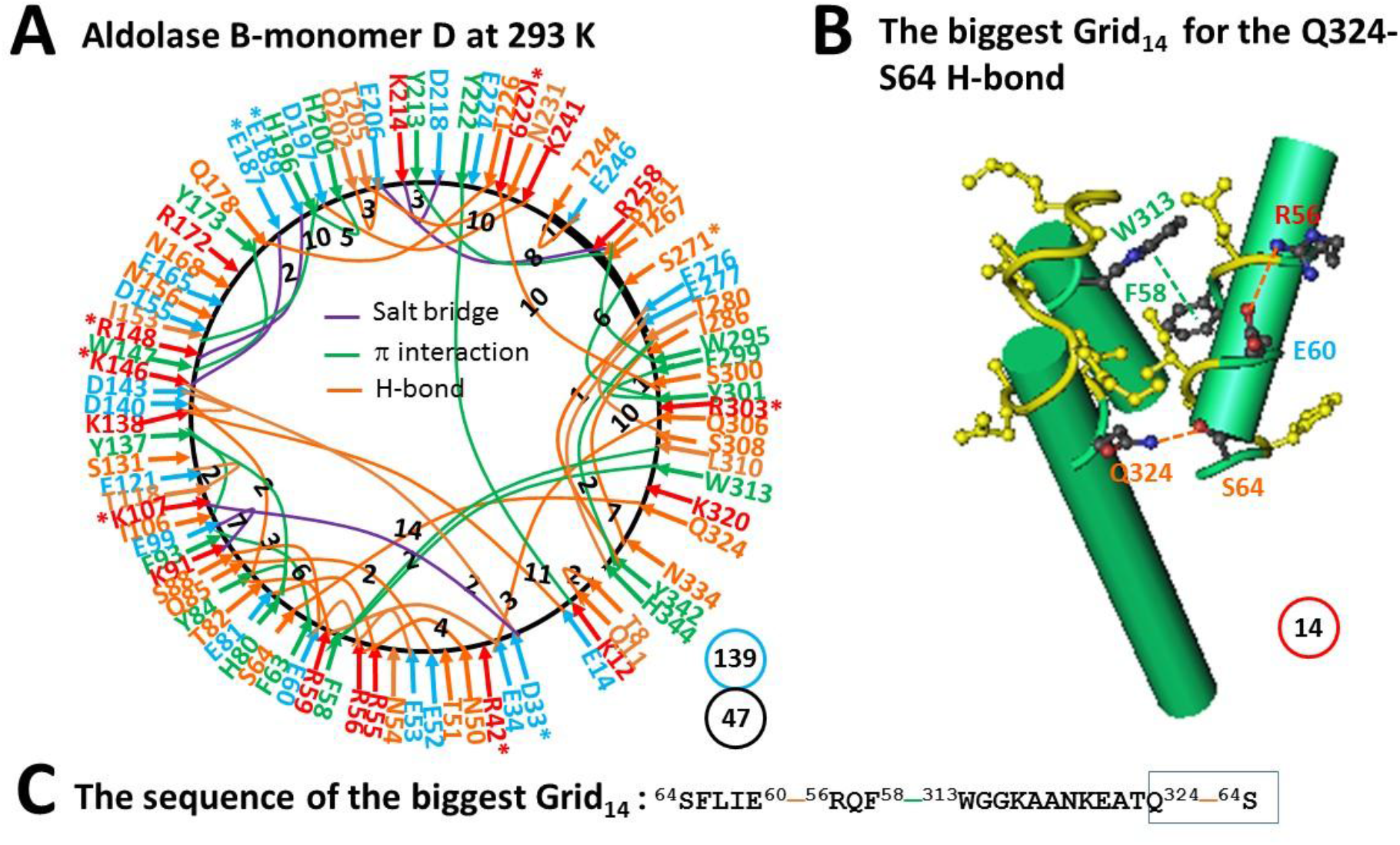
The grid-like non-covalently interacting mesh network along the single polypeptide chain of ligand-free monomer D in the aldolase B tetramer at room temperature (293 K). **A.** The topological grids in the systemic fluidic grid-like mesh network. The X-ray crystallographic structure of monomer D in aldolase B at room temperature (293 K) (PDB ID, 1QO5) was used for the model [9]. Salt bridges, π interactions, and H-bonds between pairing amino acid side chains along the single polypeptide chain from Q9 to S352 are marked in purple, green, and orange, respectively. The grid sizes required to control the relevant non-covalent interactions were calculated and labeled in black. The total grid sizes and grid size-controlled non-covalent interactions along the single polypeptide chain are shown in the blue and black circles, respectively. **B,** The structure of the biggest Grid_14_. The grid size is shown in a red circle. **C,** The sequence of the biggest Grid_14_ to control the S64-Q324 H-bond in a blue box.

### 3.6. The Biggest Grid_11_ in Monomer A of Wild-type Aldolase B at room temperature (293 K) Had a Predicted T_m_ 52 °C

Unlike monomer D of wild type aldolase B, when the N180-S131 H-bond was present again with the disrupted T82-R59 H-bond, the K146-D33 H-bond became a salt bridge, the S64-K320 H-bond came back, and the T8-Q11 H-bond shifted to the Q9-K12 one. On the other hand, the I267-W295 CH-π interaction moved to the L281-H344 one. In that case, the total grid sizes and non-covalent interactions slightly increased from 139 and 47 to 140 and 49, respectively (Fig. 6A). Thus, the systematic thermal instability was 2.86 (Table 1). When 2 equivalent H-bonds sealed the biggest Grid_11_ to control the K12-Y222 π interaction and the E14-K138 H-bond, a calculated T_m_ was about 52°C (Fig. 6B-C, Table 1).

**Figure 6.**
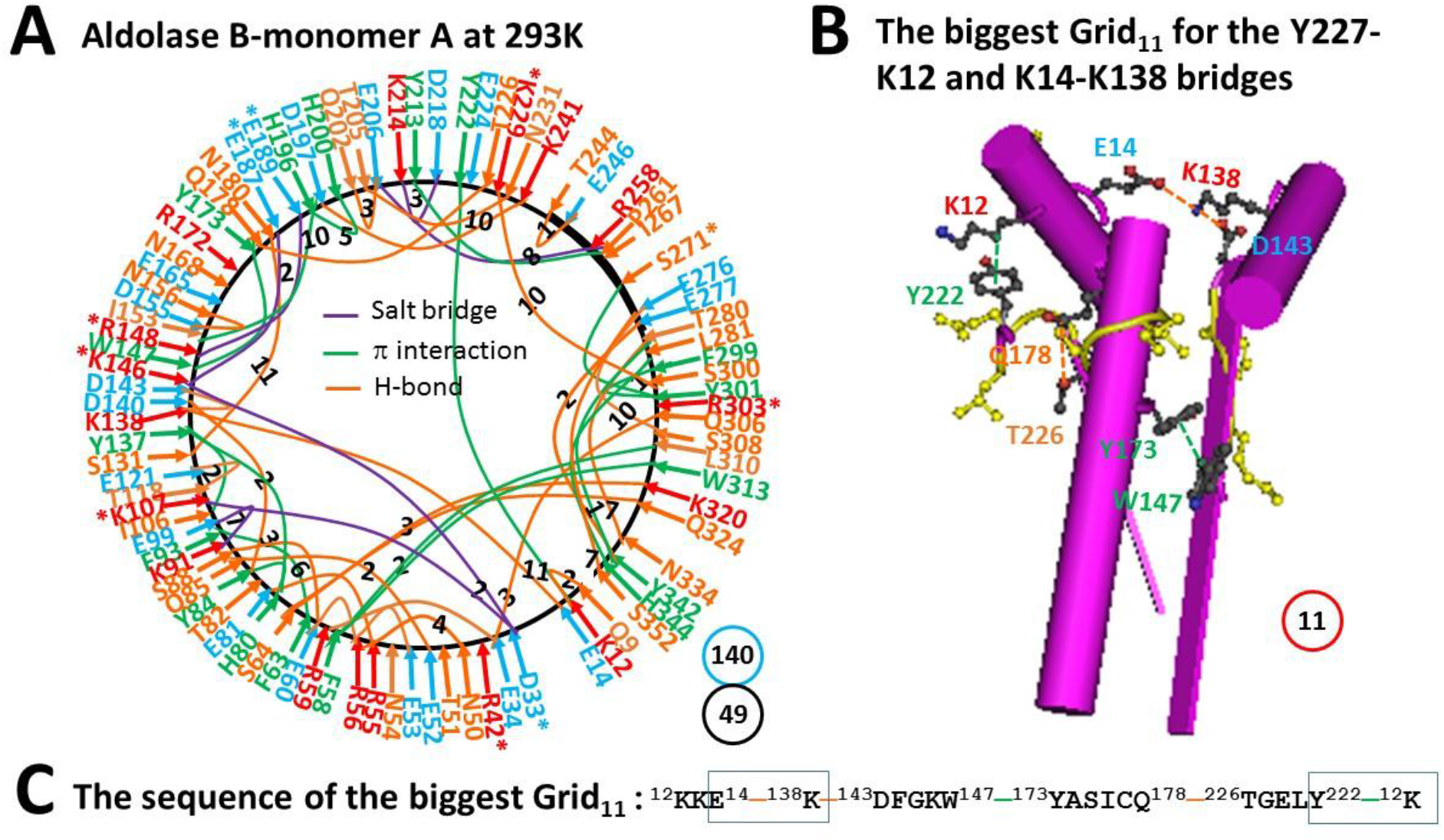
The grid-like non-covalently interacting mesh network along the single polypeptide chain of ligand-free monomer A in the aldolase B tetramer at room temperature (293 K). **A.** The topological grids in the systemic fluidic grid-like mesh network. The X-ray crystallographic structure of monomer A in aldolase B at room temperature (293 K) (PDB ID, 1QO5) was used for the model [9]. Salt bridges, π interactions, and H-bonds between pairing amino acid side chains along the single polypeptide chain from T8 to H344 are marked in purple, green, and orange, respectively. The grid sizes required to control the relevant non-covalent interactions were calculated and labeled in black. The total grid sizes and grid size-controlled non-covalent interactions along the single polypeptide chain are shown in the blue and black circles, respectively. **B,** The structure of the biggest Grid_11_. The grid size is shown in a red circle. **C,** The sequence of the biggest Grid_11_ to control the K12-Y222 CH-π interaction and the E14-K138 H-bond in blue boxes.

## 4 DISCUSSION

When non-covalent interactions such as H-bonds sealed a single polynucleotide chain to form a DNA hairpin loop, its T_m_ mainly depends on the loop size and the intensity of non-covalent interactions in the hairpin. At a given salt concentration (150 mM NaCl), a DNA hairpin with a 20-base long poly-A loop has T_th_ 34°C. When the loop size decreases from 20 to 10 bases or the H-bonding pairs in the stem increase from 2 to 4, the threshold can be increased more than 20 degrees [6]. Similarly, this computational study further proved that when non-covalent interactions such as H-bonds, salt bridges and π interactions close a single polypeptide chain to form a grid in a monomer of protein, T_m_ was primarily determined by the biggest grid.

The biggest Grid_28_ appeared in monomer B of aldolase B-A149P at 277 K. It had 2.0 equivalent H-bonds to seal a 28-residue size to control the E206-R258 H-bond so that its T_m_ was calculated as about 18 °C (Fig.1). In agreement with this prediction, this salt bridge did melt at 18 °C (Fig.2, Table 1) [2]. On the other hand, the biggest Grid_27_ was found in monomer A of the same mutant at 277 K. When 1.5 equivalent H-bond formed a 27-residue size to keep the E206-R258 salt bridge, a predicted T_m_ was about 15 °C (Fig.3). However, when the temperature increased to 291 K, the nearby Y213-P261 CH-π interaction was generated to shrink the size from 27 to 8 residues in the grid to save the salt bridge rather than disrupting it (Fig.4). Thus, the changes in the structural thermo-stability of a protein subunit at or above T_m_ could be achieved by either the disconnection of the controlled non-covalent interaction or the formation of an adjacent non-covalent interaction. In any way, the biggest grid may reduce the size to adapt a higher temperature. In the case of monomer A, the relevant size declined from 28 to 25 when the temperature raised from 277 K to 291 K (Figs. 1-2). Similarly, the relevant size in monomer B reduced from 27 to 14 (Figs 3-4). Taking together, the grid thermodynamic model could be used to predict T_m_ as T_th_ for the structural and functional changes of aldolase B and its A149P mutant.

Previous Near-UV CD studies indicated that a change in a tertiary structure of aldolase B-A149P starts at about 30°C and ends at about 40°C. In contrast, wild-type aldolase B changes the tertiary structure in a temperature range from 40°C to 48°C [1]. Because the spectral changes at 263 nm reflect the conformational alteration regarding the juxtaposition of aromatic residues [11], it is interesting to examine which biggest grid is involved in this tertiary structural rearrangement. For the aldolase B-A149P mutant, the biggest Grid_25_ had a 25-residue size to control the K12-Y222 CH-π interaction in monomer B at 277 K (Fig.2). Its T_m_ 29 °C was exactly close to the T_th_ 30 °C for the change in the tertiary structure (Table 1) [1]. Accordingly, it is proposed that the orientation of Y222 may play a critical role in the tertiary structure of the A149P mutant. Consistent with this proposal, the biggest Grid_11_ had an 11-residue size to keep the K12-Y222 CH-π interaction below the T_m_ 52 °C, which was near the end temperature 48 °C for the tertiary structural rearrangement [1]. On the other hand, the biggest Grid_14_, which was shared by monomer A of the A149P mutant at 291 K and monomer D of wild-type aldolase B at room temperature (293 K), had a 14-residue size to maintain the S64-Q324 H-bond below a T_m_ 41 °C. As this common Grid_14_ involves aromatic residues F58 and W313 and the T_m_ was close to not only the end temperature 40 °C of the tertiary structural rearrangement of the A149P mutant but also the start one 40 °C of the wild-type construct [1], these two aromatic residues may also be involved.

Prior functional studies also showed that the A149P mutant decreases the activity above 15°C until no activity was found at about 40°C while wild type aldolase B weakens the activity above 40°C until no activity was observed at about 50°C [1]. In accordance with these temperatures ranges, the biggest Grid_28_ and Grid_27_, which were observed in monomers B and A of the A149P mutant at 277 K, respectively, melted at 18 °C (Figs. 1-4). Therefore, these two biggest grids may be important for the activity of the A149P mutant (Table 1). Since the biggest Grid_14_, which was found in monomer A of the A149P mutant at 291 K and monomer D of the wild-type construct at room temperature (293 K), had a common T_m_ 41 °C (Figs. 4-5), this topological Grid_14_ may also be required for the activity of the isoenzyme (Table 1). Along with the decrease in T_m_ by the A149P mutation, the systematic thermal instability also increased from 2.86-2.89 to 3.37-3.65 (Table 1). Finally, the biggest Grid_11_ appeared in monomer A of wild-type aldolase B had T_m_ 52 °C (Fig. 6), which was close to the temperature 50 °C to terminate the activity of the wild-type construct. In this regard, this common topological Grid_11_ may be indispensable for the enzyme activity (Table 1).

Grid_11_ was highly conserved in monomer D of the wild-type construct at room temperature (293 K), monomer A of the A149P mutant at 291 K, and monomer B of the A149P mutant at 277 K. It had the shortest path from K12 to E14, K138, D143, W147, Y173, Q178, T226, Y222 and then back to K12 (Figs. 1A, 4A, 5A, 6A, 7). Thus, Grid_11_ may serve as a stable anchor for the activity of aldolase B. Although the absence of the Q178-T226 H-bond or the E187-K229 salt bridge in monomer B of the A149P mutant at 291 K may inactivate this construct at 18 °C (Figs. 1A, 2A), the replacement of the Q178-T226 H-bond by the E187-K229 salt bridge may still allow monomer A of the A149P mutant at 277 K to have a similar Grid_9_ as an anchor for the activity (Fig. 3A).

**Figure 7.**
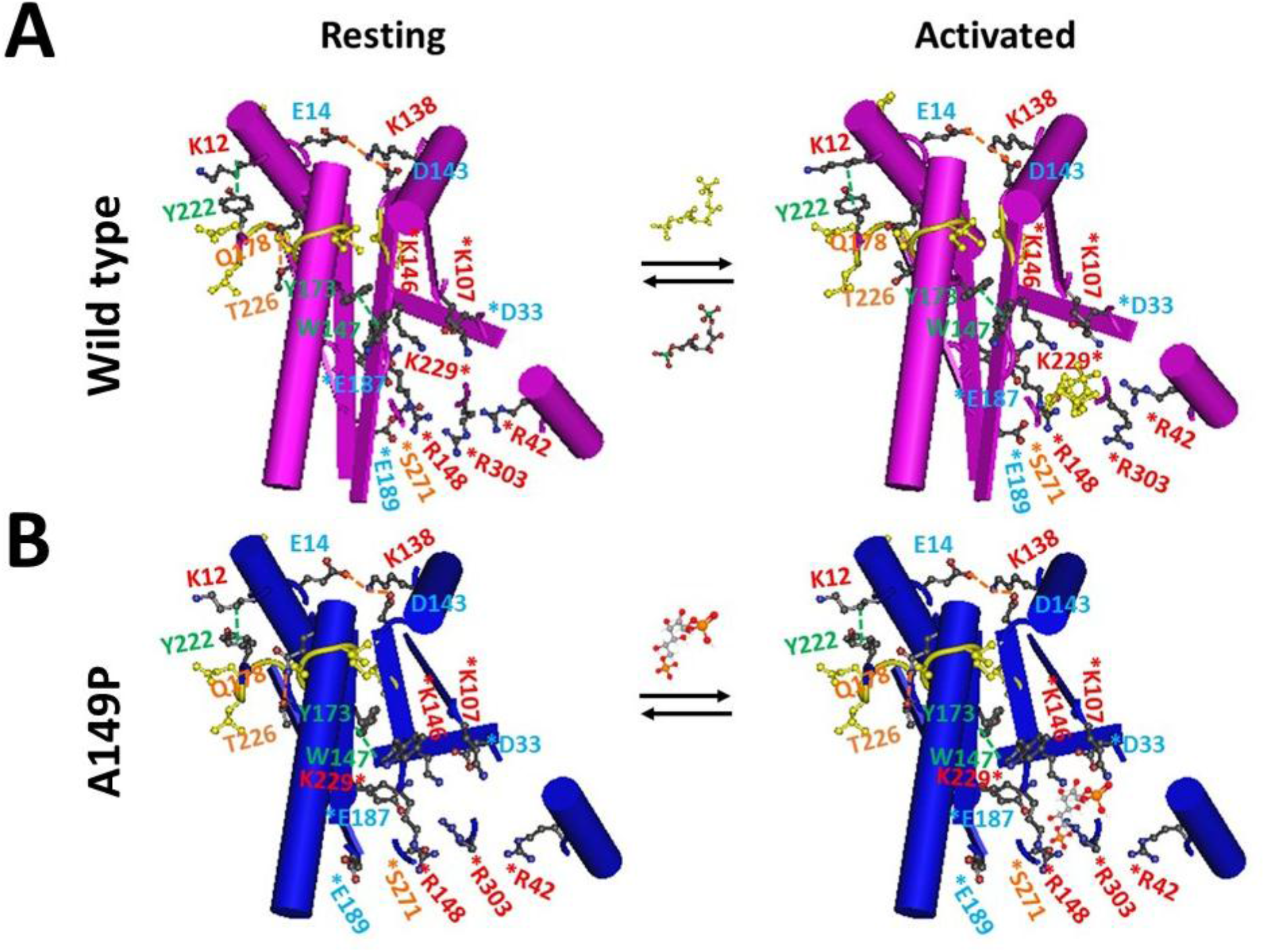
The tentative model for the activation of aldolase B and its mutant A149P upon the binding of Fru-1,6-P_2_. **A.** The X-ray crystallographic structures of aldolase B in the absence (PDB ID: 1QO5) and presence of (PDB ID: 1FDJ) Fru-1,6-P_2_. [9, 29]. **B,** The X-ray crystallographic structures of aldolase B-A149P in the absence and presence of Fru-1,6-P_2_. (PDB ID: 1XDL) [2]. A conserved and stable Grid_11_ is proposed as an anchor for active aldolase B with or without the A149P mutation. It starts from K12, goes through E14, K138, D143, W147, Y173, Q178 and T226, and ends with Y222. This Grid_11_ has at least D33, K107, K146, K148, E187, K229, S271 and R303 to secures the binding of ligands like Fru-1,6-P_2_ to the pocket.

In contrast, Aldolase B has a dynamic active site to be responsible for ligand binding, Schiff base formation, protonation of the product enamine, hydrolysis of the product Schiff base, and release of dihydroxyacetone phosphate (DHAP) from the enzyme [2, 9, 12–15]. Because this active site has residues such as D33, R42, K107, K146, R148, E187, E189, K229, S271, and R303 near Grid_11_ to realize multiple functions, both the rigid anchor and the flexible active site, which are highly conserved throughout the three aldolase isozymes A, B and C [16], may be required for the activity of this isoenzyme. Generally, E187 acts as an acid catalyst in the dehydration of the carbinolamine intermediate to support the mechanism of Schiff-base chemistry [13, 17]. The conserved R148-E189 salt bridge exposes one side of a cleft to the active site [9, 13, 18]. R303 may serve as a conformational switch in mediating the action of the C terminus during catalysis [7, 13, 19].

Since the dynamic active site is coupled to the stable Grid_11_ anchor, it is reasonable that any structural perturbation at either site by a mutation may affect the catalytic activity. In agreement with this notion, some missense K146 mutants have a marked decline in catalytic activity [20–22], the R303A/W/Q or W147R mutation affects the *k_cat_* and *K_m_* values for Fru-1,6-P_2_ [19, 23–25], the deletion of nearby residues L182 and V183 or the nearby A174D mutation in HFI patients reduces the enzyme activity [23, 26–27], and W147R and A174D are also associated with HFI [27–28]. Upon the A149P mutation, the disruption of the native β-strand structure at position 149 caused the disruption of the R148-E189 salt bridge and other structural perturbations at critical positions R148, E189 and R303 and other disordered loop regions (Figs. 3A and 4A). As a result, the orientation rearrangements of E189 and R303 enlarges the binding pocket so that the C1-phosphate-binding site of AP-aldolase is distorted in a temperature-independent manner (Fig. 7) [1–2, 9]. This loose pocket may disfavour the ligand binding so that the activity is weakened [1].

Of special note, four hydrophilic amino-acid residues form a conserved swapping H-bond network at the dimeric interfaces. They are E224 in the C-terminal β-strand from one subunit, Q202 and E206 in helix E, and R258 in helix F from the other subunit [2, 7, 9, 29]. This interfacial H-bond network may stabilize the aforementioned Grid_11_ anchor with an 11-residue size. That may be why the change in the quaternary structure also affects the enzyme activity [1].

## 5 CONCLUSION

This computational study aimed to bridge the gap between crystallographic static conformation and biochemical findings by using graph theory at an atomic level. Once the key deterministic structural factors or motifs in the systematic fluidic grid-like mesh network of non-covalent interactions were examined and identified to be responsible for a change in the tertiary or secondary structure or the specific catalytic activity in aldolase B, this study may provide a new direction to predict both the structural thermostability and the functional thermoactivity of proteins including enzymes and hence to improve the biocatalytic applications in industry.

## Conventions and Abbreviations

DHAP: dihydroxyacetone phosphate
Fru-1-P: fructose 1-phosphate
Fru-1,6-P_2_: fructose 1,6-bisphosphate
HFI: Hereditary fructose intolerance
T_m_: melting temperature
T_th_: temperature threshold

## Data Availability Statement

All data generated or analysed during this study are included in this published article.

## References

1. Malay, A. D., Procious, S. L. & Tolan, D. R. The temperature dependence of activity and structure for the most prevalent mutant aldolase B associated with hereditary fructose intolerance. Arch Biochem Biophys. 2002, 408, 295–304.

2. Malay, A. D., Allen K. N. & Tolan, D. R. Structure of the thermolabile mutant aldolase B, A149P: molecular basis of hereditary fructose intolerance. J. Mol. Biol. 2005, 347, 135–144.

3. Tolan, D. R. Molecular basis of hereditary fructose intolerance: mutations and polymorphisms in the human aldolase B gene. Hum. Mutat. 1995, 6, 210–218.

4. Baerlocher, K., Gitzelmann, R., Steinmann, B. and Gitzelmann-Cumarumsay, N. Hereditary fructose intolerance in early childhood: a major diagnostic challenge. Helv Paediat Acta, 1978, 33, 465–487.

5. Cox, T. M. Hereditary fructose intolerance. Quart J Med. 1988, 68, 585–594.

6. Jonstrup A. T., Fredsøe J., Andersen A. H. DNA Hairpins as Temperature Switches, Thermometers and Ionic Detectors. Sensors. 2013, 13, 5937–5944.

7. Dalby, A., Dauter, Z. & Littlechild, J. A. Crystal structure of human muscle aldolase complexed with fructose 1,6-bisphosphate: mechanistic implications. Protein Sci. 1999, 8, 291–297.

8. Miotto, M., Olimpieri, P. P., Rienzo, L. D., Ambrosetti, F., Corsi, P., Lepore, R., Tartaglia, G. G., Milanetti, E. Insights on protein thermal stability: a graph representation of molecular interactions. Bioinformatics. 2019, 35(15), 2569–2577.

9. Dalby, A., Tolan, D. R. & Littlechild, J. A. Crystal structure of human liver fructose 1,6-bisphos-phate aldolase. Acta Crystallog. sect. D. 2001, 57, 1526–1533.

10. Floyd, R. W. Algorithm-97 - Shortest Path. Commun Acm. 1962, 5, 345–345.

11. Schmid, F. X., in: Creighton T. E. (Ed.), Protein Structure: A Practical Approach, IRL Press, Oxford, 1989, pp. 251–285.

12. Baron, C. B., Tolan, D. R., Choi, K. H., Coburn, R. F., Aldolase, A. Ins(1,4,5)P3-binding domains as determined by site-directed mutagenesis. Biochem J. 1999, 341(Pt 3), 805–812.

13. Choi, K. H., Shi, J., Hopkins, C. E., Tolan, D. R. & Allen, K. N. Snapshots of catalysis: the structure of fructose-1,6-(bis)phosphate aldolase covalently bound to the substrate dihydroxyacetone phosphate. Biochemistry. 2001, 40, 13868–13875.

14. Arakaki, T. L., Pezza, J. A., Cronin, M. A., Hopkins, C. E., Zimmer, D. B., Tolan D. R. and Allen, K. N. Structure of human brain fructose 1,6-bisphosphate aldolase: Linking isozyme structure with function. Protein Science. 2004, 13, 3077–3084.

15. Pezza J. A., Stopa J. D., Brunyak E. M., Allen K. N. & Tolan D. R. Thermodynamic Analysis Shows Conformational Coupling/Dynamics Confers Substrate Specificity in Fructose-1,6-bisphosphate Aldolase. Biochemistry. 2007, 46, 13010–13018.

16. Lai, C. Y., Nakai, N. & Chang, D. Amino acid sequence of rabbit muscle aldolase and the structure of the active center. Science. 1974, 183, 1204–1206.

17. Maurady, A., Zdanov, A., de Moissac, D., Beaudry, D. & Sygusch, J. A conserved glutamate residue exhibits multifunctional catalytic roles in D-fructose-1,6-bisphosphate aldolases. J. Biol. Chem. 2002, 277, 9474–9483.

18. Sygusch, J., Beaudry, D. & Allaire, M. Molecular architecture of rabbit skeletal muscle aldolase at 2.7-A°resolution. Proc. Natl Acad. Sci. USA. 1987, 84, 7846–7850.

19. Choi, K. H., Mazurkie, A. S., Morris, A. J., Utheza, D., Tolan, D. R. & Allen, K. N. Structure of a fructose-1,6-bis(phosphate) aldolase liganded to its natural substrate in a cleavage-defective mutant at 2.3 A°. Biochemistry. 1999, 38, 12655–12664.

20. Morris, A. J. & Tolan, D. R. Lysine-146 of rabbit muscle aldolase is essential for cleavage and condensation of the C3–C4 bond of fructose 1,6-bis(phosphate). Biochemistry. 1994, 33, 12291–12297.

21. Morris A. J., Davenport R. C., Tolan D. R. A lysine to arginine substitution at position 146 of rabbit aldolase A shifts the rate-determining step to Schiff base formation. Protein Eng. 1996, 9, 61–67.

22. Blonski, C., Demoissac, D., Périé, J., and Sygusch, J. Inhibition of rabbit muscle aldolase by phosphorylated aromatic compounds. Biochem. J. 1997, 323, 71–77

23. Rellos P., Sygusch J., Cox T. M. Expression, purification, and characterization of natural mutants of human aldolase B: Role of quaternary structure in catalysis. J. Biol. Chem. 2000, 275, 1145–1151.

24. Santamaria, R., Tamasi, S., Del Piano, G., Sebastio, G., Andria, G., Borrone, C. Faldella, G., Izzo, P., Salvatore, F. Molecular basis of hereditary fructose intolerance in Italy: identification of two novel mutations in the aldolase B gene. J. Med.Genet. 1996, 33, 786–788.

25. Santamaria, R., Esposito, G., Vitagliano, L., Race, V., Paglionico, I., Zancan, L. Zagari, A, Salvatore, F. Functional and molecular modelling studies of two hereditary fructose intolerance-causing mutations at arginine 303 in human liver aldolase. Biochem. J. 2000, 350, 823–828.

26. Santamaria, R., Vitagliano, L., Tamasi, S., Izzo, P., Zancan, L., Zagari, A. & Salvatore, F. Novel six-nucleotide deletion in the hepatic fructose-1,6-bisphosphate aldolase gene in a patient with hereditary fructose intolerance and enzyme structure-function implications. Eur. J. Hum. Genet. 1999, 7, 409–414.

27. Cross, N. C., de Franchis, R., Sebastio, G., Dazzo, C., Tolan, D. R., Gregori, C., Odievre, M., Vidailhet, M., Romano, V., Mascali, G., Romano, C., Musumeci, S., Steinmann, B., Gitzelmann, R. & Cox, T. M. Molecular analysis of aldolase B genes in hereditary fructose intolerance. Lancet. 1990, 335, 306–309.

28. Ali, M., Cox, T. M. Diverse mutations in the aldolase B gene that underlie the prevalence of hereditary fructose intolerance. Am 7Hum Genet. 1995, 56, 1002–1005.

29. Blom, N., Sygusch, J. Product binding and role of the C-terminal region in class I D-fructose 1,6-bisphosphate aldolase. Nat Struct Biol. 1997, 4, 36–39

